# Initiation of mtDNA transcription is followed by pausing, and diverge across human cell types and during evolution

**DOI:** 10.1101/054031

**Authors:** Amit Blumberg, Edward J. Rice, Anshul Kundaje, Charles G. Danko, Dan Mishmar

## Abstract

Mitochondrial DNA (mtDNA) genes are long known to be co-transcribed in polycistrones, yet it remains impossible to study nascent mtDNA transcripts quantitatively in vivo using existing tools. To this end we used deep sequencing (GRO-seq and PRO-seq) and analyzed nascent mtDNA-encoded RNA transcripts in diverse human cell lines and metazoan organisms. Surprisingly, accurate detection of human mtDNA transcription initiation sites (TIS) in the heavy and light strands revealed a novel conserved transcription pausing site near the light strand TIS, upstream to the transcription-replication transition region. This pausing site correlated with the presence of a bacterial pausing sequence motif, yet the transcription pausing index varied quantitatively among the cell lines. Analysis of non-human organisms enabled *de novo* mtDNA sequence assembly, as well as detection of previously unknown mtDNA TIS, pausing, and transcription termination sites with unprecedented accuracy. Whereas mammals (chimpanzee, rhesus macaque, rat, and mouse) showed a human-like mtDNA transcription pattern, the invertebrate pattern (Drosophila and C. elegans) profoundly diverged. Our approach paves the path towards in vivo, quantitative, reference sequence-free analysis of mtDNA transcription in all eukaryotes.

## Introduction

Mitochondrial ATP production via the oxidative phosphorylation system (OXPHOS) is the major energy resource in eukaryotes. Because of its central role for life, OXPHOS dysfunction leads to devastating disorders, and play major role in common multifactorial diseases (Dowling 2014) such as Type 2 diabetes (Gershoni et al. 2014) and Parksinson’s disease(Coskun et al. 2012). In the vast majority of eukaryotes, OXPHOS is operated by genes encoded by two genomes – most in the nuclear genome (nDNA) and 37 in the short circular mitochondrial genome (mtDNA). This bi-genomic division is accompanied by profoundly different transcription regulatory system: whereas nDNA-encoded genes are transcribed individually by RNA polymerase 2 and the general nuclear transcription machinery, mtDNA transcription is long known to be regulated mainly by a dedicated RNA polymerase (POLRMT) and mtDNA-specific transcription factors (TFAM and TFB2) (Shutt and Shadel 2010). Moreover, mtDNA genes are co-transcribed in a strand-specific manner (Aloni and Attardi 1971): the heavy strand (i.e., 12 mRNAs, 14 tRNAs and 2 ribosomal RNAs) and light strand (one mRNA and 8 tRNAs) polycistrones, relics of the mitochondrial ancient bacterial ancestor (Zollo et al. 2012). However, as mtDNA transcription was mostly studied in vitro, little remains known about the mode and tempo of in vivo OXPHOS genes’ transcription residing on the mtDNA.

During the early 1980’s, human mtDNA transcription initiation sites were identified at a single-nucleotide resolution within the light and heavy strand promoters (LSP and HSP, respectively) (Montoya et al. 1982; Chang and Clayton 1984). These findings led to precise identification of mtDNA transcription initiation sites (TIS) in mouse (Chang and Clayton 1986a; Chang and Clayton 1986b), xenopus (Bogenhagen et al. 1986), chicken (L’Abbe et al. 1991) and in the crustacean *Artemia franciscana* (Carrodeguas and Vallejo 1997). Although such studies provided insights into the location of mtDNA promoters in the mentioned organisms, the techniques used were typically low throughput, were only semi quantitative, challenging to apply and require prior sequence knowledge. These include S1 nuclease protection and primer extension (Chang and Clayton 1984), as well as in-vitro capping (Yoza and Bogenhagen 1984). These obstacles interfered with comparative in-vivo investigation of mtDNA transcription in diverse conditions, and hampered expanding the study of mtDNA nascent transcripts to organisms lacking an mtDNA reference sequence. Finally, mtDNA transcription termination sites have been either mapped in-vitro, or were associated with MTERF binding sites (Christianson and Clayton 1986), thus, again, limiting the capability to in-vivo map transcription terminations sites in diverse organisms. It is thus imperative to develop alternative approaches.

Recently, Global and Precision-Global Run-On transcription and Sequencing assays (GRO-seq and PRO-seq, respectively) enabled high-throughput detection of nascent transcripts (Kwak et al. 2013; Core et al. 2014). Such assays can be used to resolve the genome-wide landscape of transcription start, pausing and termination sites (Kwak et al. 2013; Danko et al. 2015). In these techniques cell nuclei are isolated and run-on transcription reaction is performed in the presence of a tag that is affinity-purified to specifically isolate nascent RNA. As the cell nucleus is long known to be attached to a subset of the mitochondria (Barer et al. 1960) we reasoned that they will be co-purified with the isolated nuclei, thus potentially generating mtDNA reads. Here, we analyzed mtDNA reads generated by GRO-seq and PRO-seq experiments from 11 human cell types and seven Metazoan species. We developed a bioinformatics pipeline which identifies candidate TIS, transcription pausing and termination sites with extremely high accuracy. Such analysis revealed, for the first time, precise quantitative differences in light versus heavy strand TIS ratios between human cell types and organisms, identified candidate transcription pausing and termination sites for both the light and heavy strands in diverse organisms. Our analysis paves the path towards investigating mtDNA transcription in diverse physiological conditions, and in any given eukaryote.

## Results

### Adapting PRO-seq and GRO-seq data to analyze mtDNA transcription

GRO-seq and PRO-seq are based on massive parallel sequencing of nascent RNA extracted from either permeabilized cells or isolated cell nuclei. Since a subset of the mitochondrial population physically interact with the nuclear membrane (Barer et al. 1960), it is reasonable to assume that some of the PRO-seq reads will correspond to mtDNA transcription. To test for this possibility we analyzed GRO-seq and PRO-seq data from 11 different human cell types (Table 1, Supplementary Table S1). First, we mapped the reads using the human revised Cambridge Reference mtDNA Sequence as a scaffold (rCRS, Genbank accession number NC_012920.1). Since the human mtDNA sequence is highly variable, and hence the dense SNP map could reduce the amount of mapped reads, we used the mapped mtDNA reads to reconstruct the mtDNA sequence for each of the analyzed samples, separately. Moreover, to further increase the amount of accurately mapped reads, we took into account that the mtDNA is a circular molecule during the mapping procedure (see Materials and Methods).

**Table 1:** Mitochondrial DNA initiation and termination sites across human cell lines. Details of human GRO-seq and PRO-seq analysis.

| Cell line | Initiation |  | Light strand/<br>Heavy strand<br>Ratio | Termination |  |
| --- | --- | --- | --- | --- | --- |
|  | Light strand | Heavy strand |  | Light strand | Heavy strand |
| **AC16** | 634 | 407 | 1.88 | 16367 | 3158 |
| **CD4+** | 689 | 410 | 2.17 | 195 | 3196 |
| **GM12004** | 560 | 406 | 4.98 | 16209 | 3154 |
| **GM12750** | 560 | 407 | 2.41 | 16182 | 3163 |
| **GM12878** | 562 | 407 | 3.14 | 16205 | 3144 |
| **Hela** | 560 | 408 | 0.87 | 16268 | 2612 |
| **IMR90** | 561 | 407 | 0.63 | 16259 | 3252 |
| **Jurkat** | 675 | 410 | 2.35 | 16076 | 3191 |
| **K562** | 658-662 | 407-409 | 1.47 | 16209-16403 | 2712-3238 |
| **MCF7** | 561 | 408 | 8.77 | 16170 | 3017 |
| **U20S** | 560 | 407 | 1.38 | 16288 | 2855-3201 |

### Analysis of NUMTs confirmed mtDNA mapping specificity

The isolation of cell nuclei during sample preparation for both GRO-seq and PRO-seq raised the possibility that a subset of the identified mtDNA reads reflect contamination by mtDNA-like pseudogenes that have been transferred to the cell nucleus during the course of evolution (NUMTs) (Hazkani-Covo et al. 2003; Mishmar et al. 2004). To control for this possibility we focused our analyses on regions encompassing the light and heavy strand mtDNA promoters (LSP and HSP, respectively). As GRO-seq and PRO-seq data sequence cDNA generated from nascent RNA extracts, we first BLAST searched our identified mtDNA reads against the entire *Homo Sapiens* Ref Seq RNA database (https://blast.ncbi.nlm.nih.gov/Blast.cgi). To increase sensitivity of our NUMT screen, while taking into account the relatively short read length generated by GRO-seq and PRO-seq (i.e. a minimum of 30 bases), we focused our screen on nuclear genomic BLAST hits that were longer than 28 bp. This screen did not reveal any candidate NUMTs that mapped within the promoters region of both mtDNA strands (Supplementary Fig. S1). To increase our stringency, we expanded our BLAST analysis to DNA reads of the entire human genome (GRCh38). This screen revealed three BLAST hits: one from chromosome 5, within the region spanning the LSP (hereby referred as the light strand NUMT), and additional two BLAST hits from chromosome 5 and chromosome 11, respectively, that span the HSP (hereby referred as heavy strand NUMT 1 and 2). The light strand NUMT diverged from the rCRS in three mtDNA positions (i.e. 369, 377,401); the heavy strand NUMTs diverged from the rCRS in eight mtDNA positions (i.e., 572, 573, 576, 592, 596, 686, 710, 711). Analysis of the DNA sequences in mtDNA mapped reads indicated that only 0.21% of the reads encompassing the LSP could be explained by NUMT contamination (SD=0.007). Similarly, only 0.6% of the heavy strand reads corresponded to candidate NUMTs in the region encompassing HSP1 (SD=0.017) and only 0.013% of the reads (SD=0.047) corresponded to NUMTs within the region encompassing HSP2 (Table 2, Supplementary Table S2). Since the proportion of NUMT reads was very low, and hence is expected to have only negligible impact on our transcription analysis, we avoided unique mtDNA mapping in further analyses.

**Table 2:** Identification of NUMts in all human cells lines tested. Identification of NUMTs within human mtDNA TIS.

|  | LSP |  |  |  | HSP1 |  |  |  | HSP2 |  |  |
| --- | --- | --- | --- | --- | --- | --- | --- | --- | --- | --- | --- |
| Position | 368 | 377 | 401 | 572 | 573 | 576 | 592 | 596 | 686 | 710 | 711 |
| mtDNA nucleotide | A | C | T | C | C | A | C | T | A | T | T |
| NUMT nucleotide | G | T | C | A | T | G | T | C | G | C | C |
| Range of mtDNA reads | 89-100021 | 83-101386 | 65-77091 | 10- 699 | 11-707 | 12-747 | 20-747 | 21-736 | 7-2125 | 5-3123 | 5-3193 |
| Range of %NUMTs reads | 0%-4.47% | 0%-0.05% | 0%-0.12% | 0%- 10% | 0%-1.27% | 0 | 0%-2.9% | 0% | 0%-0.19% | 0% | 0% |
| Average of %NUMTs reads (SD) | 0.58% (1.18%) | 0.01% (0.02%) | 0.04% (0.04%) | 1.3% (3.29%) | 0.23% (0.44%) | 0% (0%) | 1.49% (1.15%) | 0% (0%) | 0.04% (0.08%) | 0% (0%) | 0% (0%) |

### Identification of mtDNA transcription initiation sites at the light and heavy strands in diverse human cell types

Having shown that PRO-seq and GRO-seq can be used to analyze mtDNA transcription, we next sought to identify candidate transcription initiation sites (TIS). We screened mtDNA for regions harboring no mapped reads followed by sudden increase in downstream reads (Fig. 1). To increase our sensitivity we used a two-step approach (Fig. 1). The first step (step 1) was aimed towards crude identification of the best candidate TIS: we normalized the sequence coverage of each nucleotide position to the average in sliding 200 bp windows. Next, we searched for mtDNA nucleotide positions with the following characteristics: upstream (200-1000 bases window) read coverage of less than 5% of the average read coverage across the entire mtDNA in combination with downstream (500 bases) read coverage greater than 5% of the mtDNA average read coverage. In samples lacking nucleotide positions that passed these criteria, the read coverage threshold was increased by 1% increments, until such positions were identified. The score for each of these sites was the ratio of downstream (50 bases) to upstream (500 bases) read coverage. Notably, if the distance between the two positions was greater than 1Kb we divided them into separate units. Finally, scores were calculated for the candidate TIS of each transcription unit (if there were more than one). The second analysis step (step 2) was employed to sort for the best TIS among the candidates identified in step 1. To this end we re-analyzed the read coverage per nucleotide, and re-calculated the downstream (50 bases) versus upstream (250 bases) ratio for positions +/- 100 nucleotides relative to the candidate positions listed in step 1. The nucleotide position with the highest score served as the best candidate TIS.

**Figure 1.**
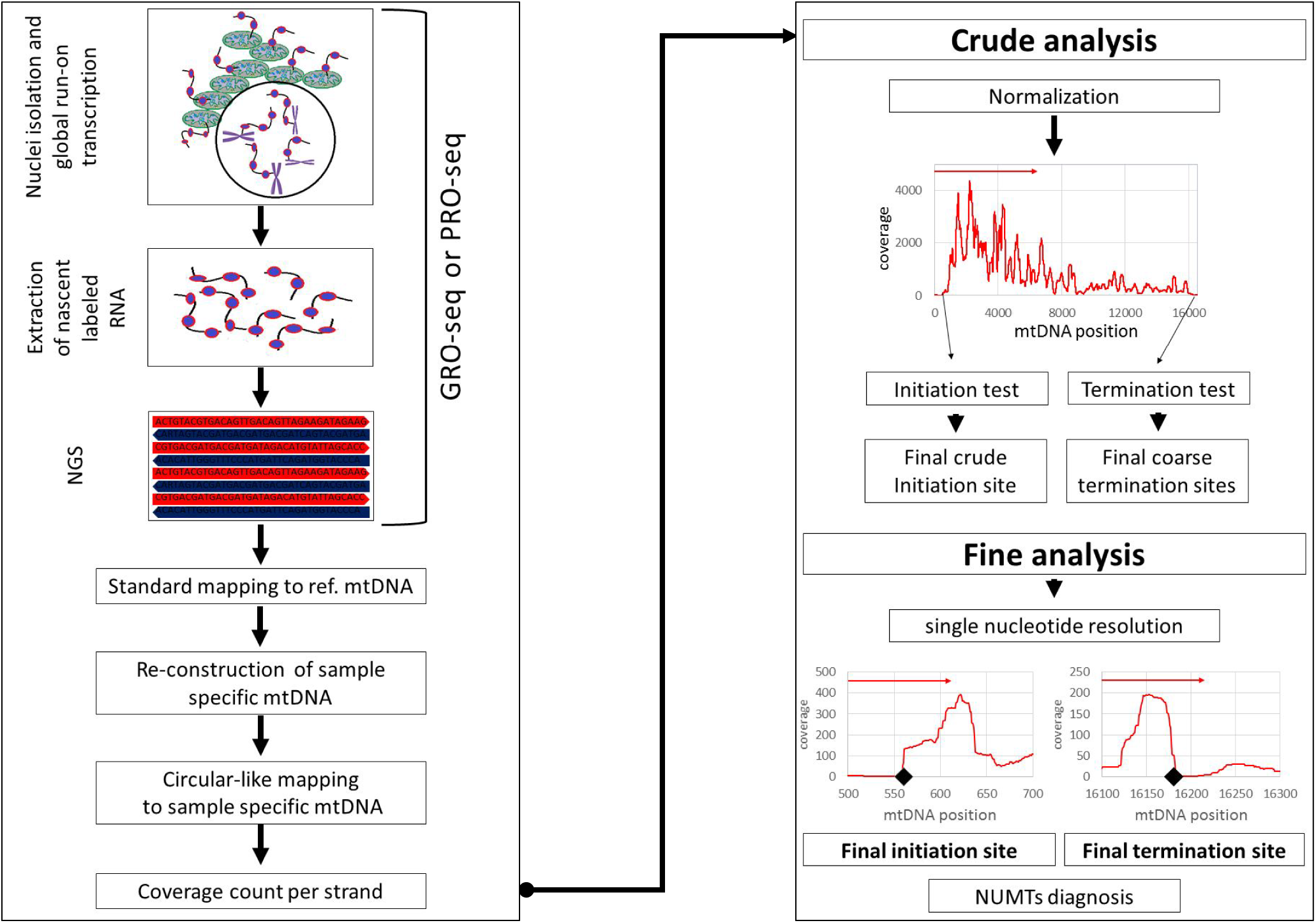
Workflow of analysis: A. GRO-seq and PRO-seq experiments generate genome-wide nascent transcript data. Although these techniques are based on nuclei isolation, mitochondrial DNA (mtDNA)-encoded RNA reads are generated since some mitochondria are attached to the cell nucleus. The extracted mtDNA sequences enable reconstruction of sample-specific mtDNA sequence which is used, in turn, as a circular-like mapping reference. This allows counting the sequencing read coverage in a strand-specific manner. B. Analysis of mtDNA transcription initiation and termination sites. Two steps were designed – (I) a crude step for candidate transcription initiation site (TIS) identification– identifying abrupt increase in read coverage in a nucleotide resolution within 200 bp sliding windows. (II) Fine analysis – focusing on highest scoring regions to identify the nest TIS candidate. Notably, the identification of transcription termination sites utilizes the same approach, yet instead of abrupt increase in reads – abrupt decrease in read coverage is sought for.

Firstly, we applied our approach to analyze PRO-seq data from K562 cells. Our analysis indicated that the TIS of both human mtDNA strands were perfectly consistent with known mtDNA promoters (Table 1). Specifically, consistent with previous findings, the TIS at positions 409 was exactly within the known LSP, and the major heavy strand TIS was within positions 658, right downstream to position 645, exactly within the identified heavy strand promoter 2 (HSP2) (Zollo et al. 2012). Secondly, previous estimates of higher transcription signal intensity near the promoter of the light strand (Chang and Clayton 1984) were corroborated using PRO-seq, i.e. the read density of the entire light strand was 1.63 fold higher than the read density of the entire heavy strand). Notably, while employing unique mtDNA mapping against the entire human genome (hg19) the region encompassing the light strand TIS was precisely identified, yet several candidate heavy strand TIS emerged, thus preventing precise identification of the best TIS candidate. Hence, unique mapping did not improve our precision. Finally, to control for possible methodological differences between the GRO-seq and PRO-seq assays, we identified TIS separately in available GRO-seq and PRO-seq experiments from the K562 cell line, and identified identical TIS predictions.

### mtDNA mode of transcription diverge across human cell types

Encouraged by our precise TIS identification in K562 cells we tested for possible variability of mtDNA transcription among human cell types. Since we analyzed primary transcription, read coverage is expected to differ across the mtDNA sequence. Sequencing read coverage not only diverged across the mtDNA but also varied among 11 tested cell types (Supplementary Table S3). For the entire heavy strand the range of total coverage per nucleotide position was 2.71-14900 (mean=1395.69, SD=3431.15) and the number of positions with coverage greater than 1% of the total sequence coverage ranged between 6822 to 16402 in the different cell lines (mean=14777.94, SD=2371.39). Considering the entire light strand, the range of total coverage was 24.21-24317.09 (mean=2453.07, SD=5581.18) and the number of positions with coverage greater than 1% of the total coverage ranged between 12623 and 15191 between the cell lines (mean=14191.88, SD=629.78). As such differences were consistent between experimental replicates, our results suggest profound quantitative variation in mtDNA transcription levels among human tissues.

With this in mind, we asked whether we could identify differences in the ratio of reads mapping to the HSP or LSP between cell lines. Our analysis of the tested human cell lines uncovered a surprising amount of variability in the location of the TIS between different cell lines. Firstly, similar to K562 cells, we found that the light strand TIS of all samples was located within mtDNA nucleotide positions 406-410, in perfect match with the known human LSP. Seven out of the 11 tested cell lines revealed a heavy strand mtDNA TIS within nucleotide positions 560-562, exactly within the known HSP1. Samples with very high read coverage tended to provide highly reproducible results in duplicate experiments as can be seen in U20S cells. Considering the other four samples, heavy strand TIS was located within positions 634 and 689, downstream of the known HSP1 and closer to HSP2. Notably, patterns did not vary between experiments that used isolated cell nuclei or whole permeabilized cells, thus partially controlling for possible over-representation of certain subcellular mitochondrial population.

Differences between the read density in the heavy and light strands are also consistent with previously shown higher activation of the light strand as compared to heavy strand promoters(Chang and Clayton 1984) in most of the tested cells (9 out of 11). Nevertheless, while calculating the read density of the entire light strand and heavy strand, respectively, such ratios differed among cell types (Fig. 2, Table 1). Specifically, the highest light strand/heavy strand transcription ratio (~9 fold) was calculated for MCF7 cells, and the lowest light strand/heavy strand read density (~0.6) was calculated for IMR90 cells. For the remaining nine cell lines, the calculated light strand/heavy strand read density ratios ranged between 1.3 to 5 fold. This suggests, for the first time, profound quantitative variation in mtDNA transcription initiation patterns among human tissues

**Figure 2.**
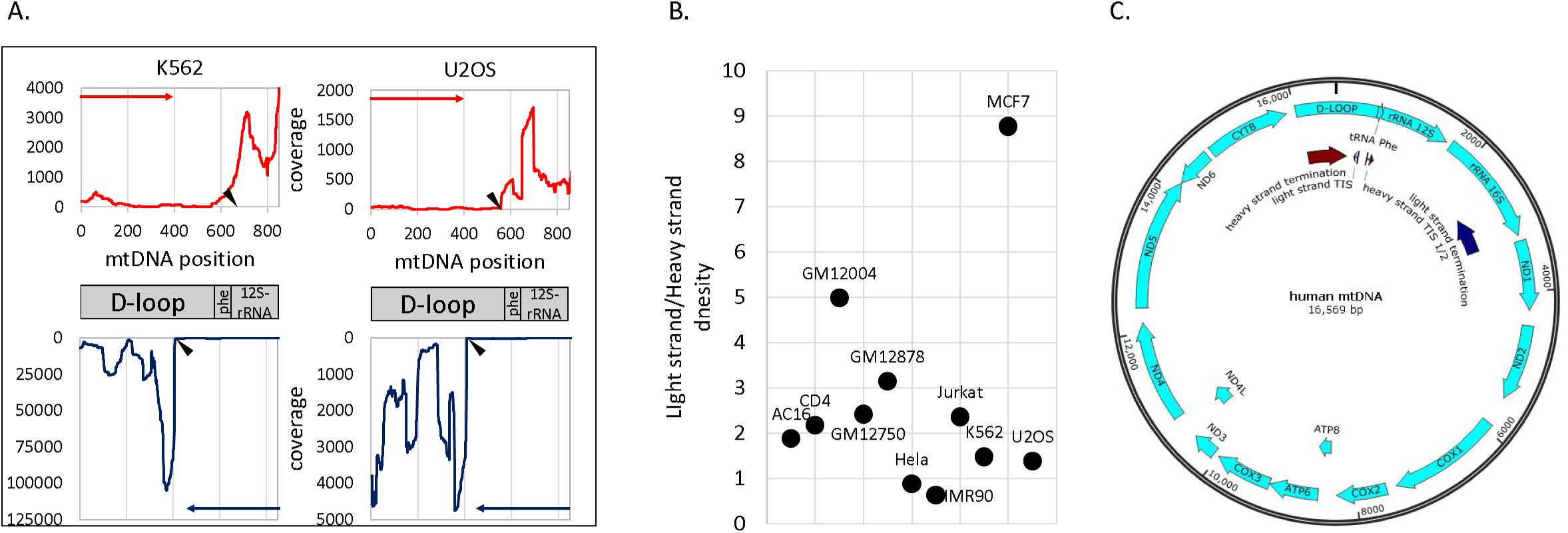
Accurate identification of human mtDNA TIS: A. Sequencing reads coverage around the mtDNA TIS in two cell lines. Upper panel – sequencing read coverage pattern of the mtDNA heavy strand (red); lower panel - sequencing read coverage of the light strand (blue). Putative TIS is designated by black triangle. Y-axis: sequencing read coverage. X-axis: mtDNA position. Left panel – PRO-seq experiment of the K562 cell line. Right panel - GRO-seq experiment from the U2OS cell line. B. Ratio of sequencing coverage of 100 bp downstream to the light strand TIS (LST) and the coverage of 100 bp downstream to the major heavy strand TIS (HST). Y-axis: LST/HST ratio. X-axis: human cell type (in brackets-mtDNA haplogroup). C. Summary of mtDNA transcription pattern - PRO-seq and GRO-seq experiments in 11 human cell types: **TIS** - Light strand TIS was identified in all tested human cell types (N=11) in positions 407-410. In most of the tested cells (7 of 11), the major heavy strand TIS was mapped in positions 560-562 (TIS 1).In the remaining cell types (4) the major heavy strand TIS was located in positions 634-689 (TIS 2). **Termination sites** – Light strand transcription termination was identified within the 16S rRNA gene, in the region encompassing positions 2612-3252 (dark blue arrow). Heavy strand transcription termination was identified within the D-LOOP, between positions 16076-195 (dark red arrow).

### Transcription pausing occurs immediately downstream to the Light strand TIS

The sequencing read pattern encompassing the light strand TIS appeared very different from that of the heavy strand TIS. Unexpectedly, in the light strand we observed a sharp peak of read coverage immediately downstream (~50 nucleotide distance) to the TIS, whereas the read pattern right downstream of the heavy strand TIS appears to be ragged (Fig. 2). Such PRO-seq peak pattern in the nuclear genome was previously interpreted as pausing sites of the transcription machinery (Kwak et al. 2013) [see below]. To determine whether a detectible enrichment of transcriptionally competent RNA polymerase was found in either HSP or LSP we assessed the read coverage in 10 nucleotide sliding window downstream to the TIS in all tested human cell lines, and found a significant enrichment of reads near in the light strand TIS, which was consistently located in positions 356-380, 30-50 nucleotides from the TIS. In the heavy strand we found that only in 2 out of 11 cell lines there were significantly enriched pausing peaks: in MCF7 and IMR90 cells the candidate pausing peaks were located at positions 597 and 653, respectively. These results were consistent between GRO-seq and PRO-seq experiments.

To determine the precise coordinates of the mitochondrial RNA polymerase near the transcription active site we focused further analysis on PRO-seq experiments, which are designed to resolve RNA polymerase progression at a single nucleotide resolution, in three cell lines (K562, Jurkat, and CD4+ T-cells). We mapped the position of each read using only the precise coordinates of the 3’ end. We found that the light strand pause site occurred within mtDNA positions 355-361. Since the peak morphology of the heavy strand was ragged, and the pausing index was very low (<0.1), mapping the candidate pausing site was less accurate (mtDNA positions 677-715). Secondly, such mapping was limited only to the K562 cell line, since Jurkat and CD4+ cells had lower sequence coverage at these positions (less than 10X) and hence were less informative. We interpret these results to mean little or no transcription pausing near the heavy strand TIS, but a robust pause on the light strand TIS. Notably, the pausing index in the light strand varied ~44 fold between the tested cell lines (Fig. 3 and Supplementary Table S4). This suggested quantitative tissue-specific mtDNA transcription pausing differences.

**Figure 3.**
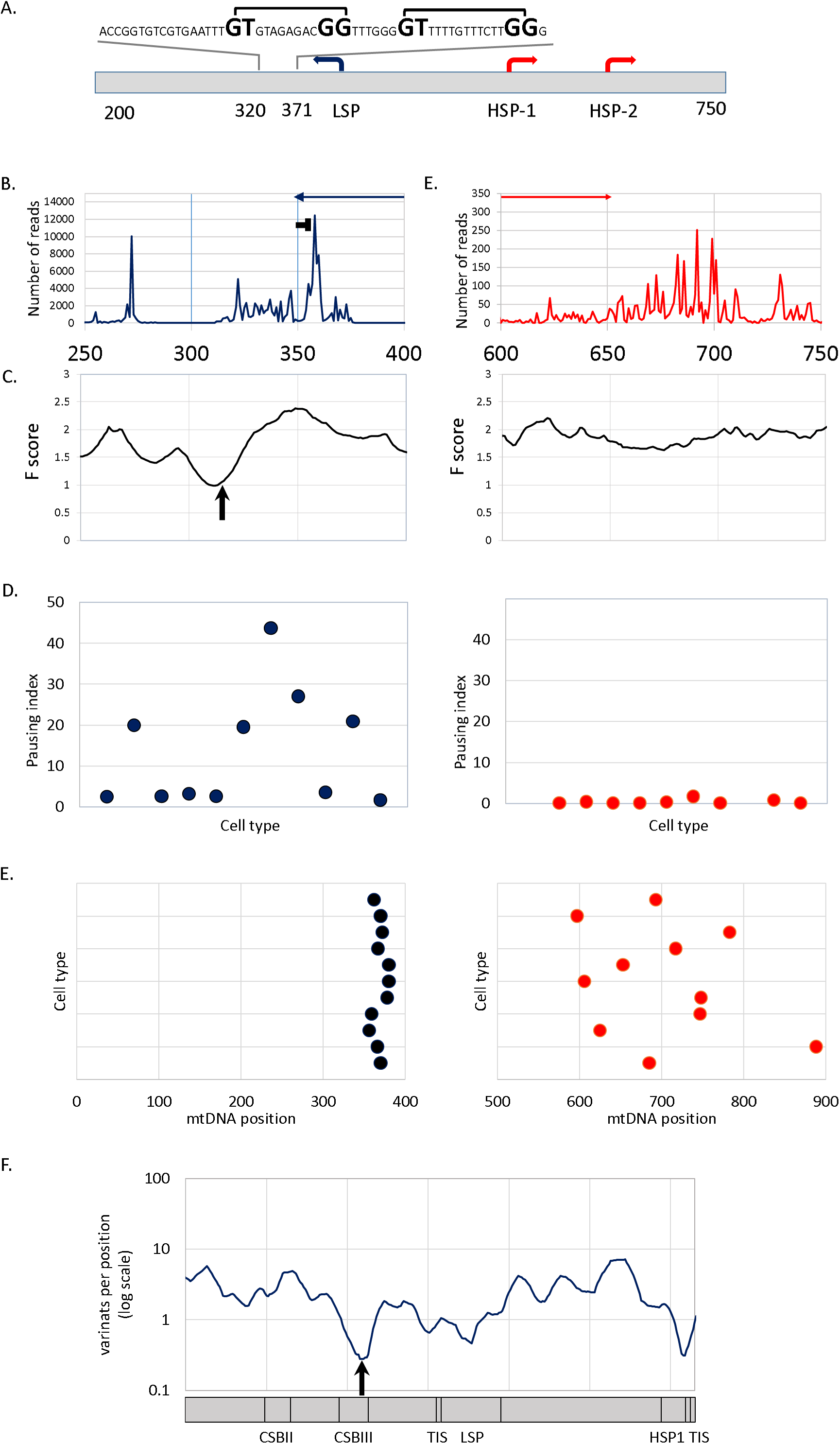
mtDNA Transcription consistently pause at distinct sites near the heavy and light strand TIS: A. mtDNA transcriptional regulation elements. Presented is the complementary human sequence of the light mtDNA strand. The mtDNA sequence around the pausing peak is above the illustrated graph; square bracket - the bacterial pausing motif. The mandatory nucleotides within the motif are highlighted by larger font size. B. Coverage of the 3 prime end of the PRO-seq experiment from K562 cell line. X-axis: mtDNA nucleotide position. Y-axis: the upper panel – number of reads in the 3 prime. Colored arrows (blue and red) - the direction of the light and heavy strand transcription, respectively. The ‘horizontal T’ sign represent the pausing site. C. DNase-seq experiment from K562 cell line. X-axis: mtDNA nucleotide position. Y-axis: F score of DNase-seq analysis (see materials and methods). The lower the score the more protected is the DNA by proteins. The black arrow point to the DHSS. D. Pausing index across human cell types. Left panel – light strand, right panel – heavy strand. X-axis - human cells type, Y-axis – pausing index values. E. Pausing site nucleotide position across human cell types. Left panel – light strand, right panel – heavy strand. X-axis – mtDNA position, Y-axis – human cells type. F. Human population SNPs density. X-axis – mtDNA position, Y-axis – SNPs density measured as variants per position (log scale).

We next sought to address the mechanism that underlies pausing near the light strand TIS. Although Metazoan systems establish nDNA transcription pausing by specific protein complexes, including DSIF and NELF (Kwak and Lis 2013), these protein complex are strictly nuclear localized and have not been characterized in the mitochondria. As mitochondria originated from an ancient bacterial symbiont we hypothesized that the transcription pausing sites may harbor bacterial-like attributes. To test this hypothesis we searched for the presence of a ~15bp sequence motif responsible for transcription pausing in *Escherichia coli* by destabilizing bacterial RNA polymerase (Larson et al. 2014; Vvedenskaya et al. 2014). Strikingly, we found two such tandem motifs within the light strand pausing peak (Fig. 3). As an alternative model, we also tested whether pausing occurs because POLRMT encounters a DNA bound protein. We analyzed available DNase-seq data for six cell lines. This analysis revealed a DNase hyper sensitivity site (DHSS) right downstream to the light-strand TIS, suggesting that an mtDNA-bound protein is involved in the light strand transcription pausing. In contrast, no DHSS was identified downstream to the heavy strand TIS, which also lacked a pausing site as mentioned above.

We next sought to assess the functional importance of mutation in the TIS at both strands, as well as that of the light strand pausing site. Since genome editing technology has yet to be established for the mtDNA in cells, alternative approaches should be used to assess the functional importance of mtDNA sequences. Hence, to assess whether the sequence motif putatively responsible for paused polymerase is important for mtDNA genome function, we asked whether DNA sequence encoding the pausing motif is conserved during the course of evolution. We studied SNP density in humans at the pausing site and TIS, as a proxy for signatures of selection. We found that the frequency of mutational events within the light strand pausing site, was significantly lower than the rest of the D-Loop (Fig. 3; positions 358-360, 0.28 variants per position – normalized to 10 base windows; p=0.037). Similarly, the frequency of mutational events around HSP1 was also significantly lower (positions 558-559, 0.31 variants per position – normalized to 10 bases; p=0.045), although the reduced frequency of mutational events at the LSP was only marginally significant (position 427, 0.46 variants per position – normalized to 10 bases; p= 0.072). These results imply that in vivo mtDNA transcription pausing at the light strand is not only common to all cell lines tested, but is also negatively selected and hence is likely to be functionally significant.

### Identification of mtDNA transcription termination sites

To date, mtDNA transcription termination sites have been determined in vitro (Christianson and Clayton 1986), and were correlated with the mtDNA binding sites of the transcription termination factors of the mTERF family (Park et al. 2007). Given our successful precise identification of TIS in both mtDNA strands, we attempted to identify candidate mtDNA transcription termination sites. To this end we employed the similar set of criteria as those applied while identifying TIS, yet instead of looking for regions in which the sequencing read coverage was dramatically increased, we looked for the opposite – regions in which the read coverage had dramatically dropped. We found that mapping heavy and light stands termination sites was less precise than TIS, although all tested samples revealed candidate termination sites within the same regions (Fig. 2). This argues for gradual rather than an abrupt transcription termination process. Specifically, we found that transcription termination of the light strand occurred between mtDNA positions 2619 and 3259, corresponding to the 3` end of the 16S rRNA gene. The heavy strand termination was identified between positions 16056 and 194 within the D-loop in all cell lines tested.

### Human mtDNA RNA-DNA differences

GRO-seq and PRO-seq experiments were recently used to estimate the timing at which RNA-DNA Differences (RDDs) occur during transcription of nDNA genes (Wang et al. 2014). Recently, we found A-to-U and A-to-G RDDs in human mtDNA position 2617 (Bar-Yaacov et al. 2013), and were curious whether they appeared already at the early stages of transcription. Analysis of all human GRO-seq and PRO-seq data available to us revealed that the RDDs were represented by less than 1% of reads encompassing mtDNA position 2617 in all samples (Supplementary Table S5) as opposed to more than 40% of the steady state mitochondrial transcripts (Bar-Yaacov et al. 2013). Hence our data is consistent with likely post transcriptional accumulation of the 2617 RDD in humans.

### Identification of mtDNA transcription initiation and termination sites in divergent Metazoans

Our successful identification of transcription initiation and termination sites in humans urged us to test our approach on non-human organisms, lacking previous experimental data and accurate mtDNA TIS mapping. As the first step, we analyzed available PRO-seq data generated by us and others from CD4+ lymphocytes from mammals (i.e., chimpanzee, rhesus macaque, rat and mouse). We found that the general mammalian pattern of mtDNA transcription initiation and termination was quite similar to humans. Specifically, in chimpanzee, rat and mouse transcription initiation, termination and pausing exhibited similar pattern to humans (Fig. 4A): a distinct pausing peak 28-36 bases downstream to the light strand TIS, a light strand transcription termination around the 3’ end of 16S rRNA gene, and a heavy strand TIS (Fig. 4B). In Rhesus Macaque, the pattern was somewhat different: the pausing site was more than 100 bases downstream to the light strand TIS (Fig. 4B). The mtDNA TIS and termination pattern of the heavy strand in the chimpanzee, rat and mouse was generally similar to that of humans, with transcription initiation occurring right upstream to the tRNA^−phe^ gene (corresponding to the putative HSP1) and termination within the D-loop. The Rhesus Macaque heavy strand TIS mapped downstream to tRNA^−phe^ and likely correspond to HSP2. The Light strand/Heavy strand TIS ratio (calculated as described for human samples) varied among species (Supplementary Table S6). Moreover, the ratio between the overall read coverage across the coding regions of the light and heavy strands notably varied among species: While the read coverage of the mtDNA heavy strand was twice the coverage of the light strand in rat and mouse, this ratio became nearly one to one in Rhesus. In humans and chimpanzee an opposite pattern emerged, with 2 fold higher coverage of the light strand as compared to the heavy strand coding regions (Supplementary Table S6).

**Figure 4.**
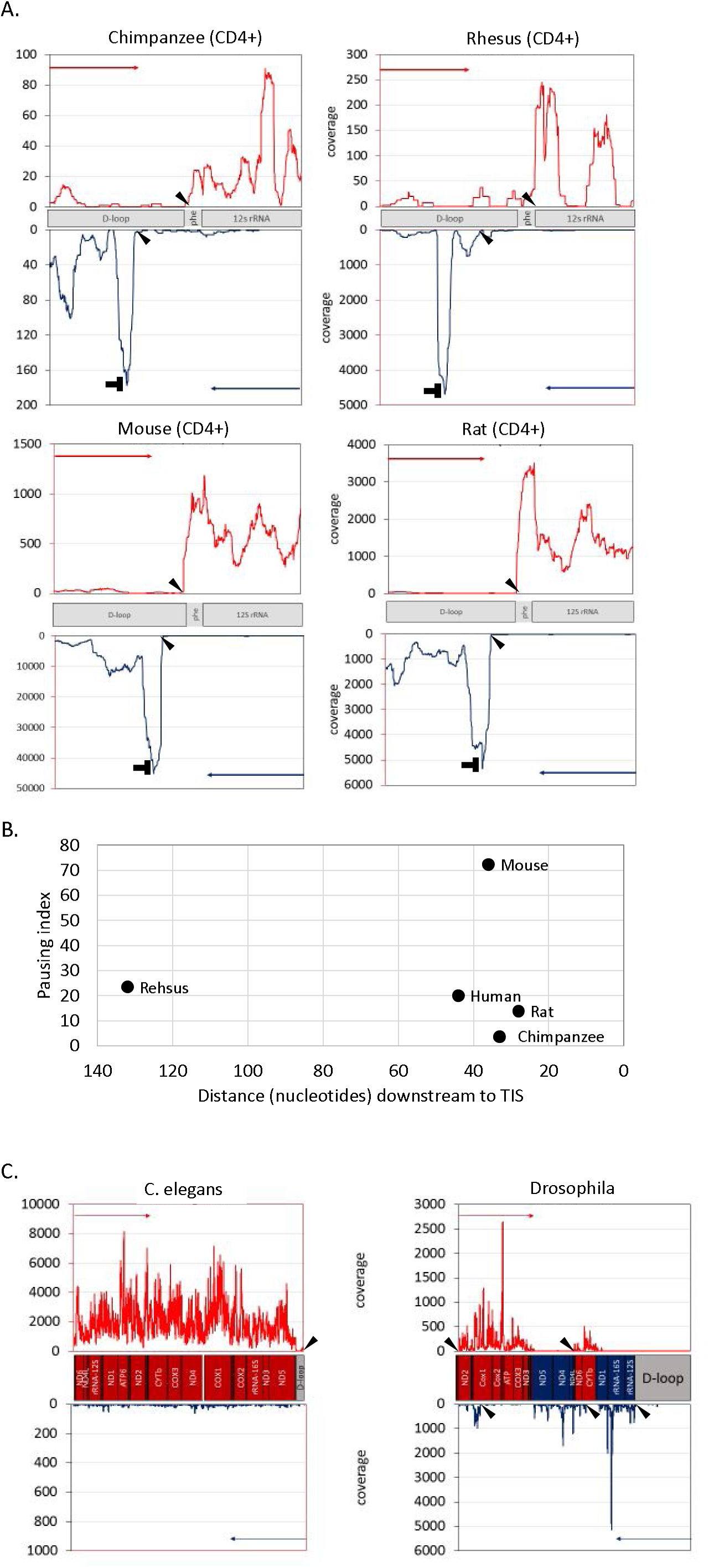
Identification of mtDNA nascent transcript across evolution: A. PRO-seq experiment performed in 4 mammalian CD4+ cells: Chimpanzee, Rhesus, Rat and Mouse. X and Y axis – are identical to Figure 2, and ‘horizontal T’ sign designate the pausing site. Arrowheads in all panel point to the calculated identified TIS. Notably, in three species (chimpanzee, rat and mouse) the major heavy strand TIS was identified downstream to the tRNA phenylalanine gene, similar to the human heavy strand TIS 1. B. Pausing site of light strand transcription in mammals. X-axis–distance (in nucleotides) of the pausing site from the light strand TIS. Y-axis – pausing index value of each species. The name of each species is indicated to the right of each dot. C. Left panel – analysis of GRO-seq data from C. elegans. In this species there is a single TIS for a single transcription unit, present only at the heavy strand. Right panel – analysis of PRO-seq data from Drosophila. Five candidate TIS were identified – two in the heavy strand and three in the light strand.

We next employed our approach to identify mtDNA TIS and termination sites in invertebrates - Drosophila and *C. elegans*. Our analysis revealed a single mtDNA TIS in *C. elegans*, only in the heavy strand (Fig. 4C), in perfect match with gene content: in the worm, all genes are encoded by the heavy mtDNA strand. In Drosophila, in consistence with previously observed transcription units using RNA-seq analysis (Torres et al. 2009), our nascent RNA analysis revealed two TIS for the heavy strand, three major and two minor candidate TIS for the light strand (Fig. 4C). Similar to *C. elegans*, the Drosophila pattern exactly corresponded to gene content. Notably, the two minor light strand TIS (one located at mtDNA position 17110 within the control region, and the other at position 3012 within the first heavy strand transcription unit – according to GenBank accession NC_024511.2) did not correspond to any of the previously described transcription units in Drosophila.

### GRO-seq and PRO-seq data enabled de-novo assembly of the mtDNA in non-human organisms

Since the mtDNA is present in high copy number across all studied eukaryotes, sequence coverage is expected to be sufficiently high to enable de-novo mtDNA sequence assembly. As this might be very useful for organisms lacking a reference genome, we assessed our capability to de-novo assemble the mtDNA sequence using GRO-seq and PRO-seq data, from Drosophila *and C. elegans* as test cases. For the assembly of the mtDNA in both Drosophila and *C. elegans* we used the mtDNA of phylogenetically-related species as a scaffold (*Bactrocera arecae* and *Litoditis aff. Marina Pml*, respectively). Drosophila mtDNA sequence contigs encompassed 80.4% of the used mtDNA scaffold, in comparison to 86.8% coverage, when the species-related scaffold was replaced by the known Drosophila mtDNA reference sequence. In *C. elegans* the reconstructed sequence contigs encompassed most (97%) of the species-related scaffold, as compared to 98.3% coverage, when we used the known *C. elegans* mtDNA as a reference sequence (Supplementary Fig. S2). The gaps in both studied species mostly corresponded to non-coding mtDNA regions, which are known to vary in length among species (Supplementary Fig. S2).

## Discussion

Here, we analyzed modes of early mtDNA transcription in diverse cell lines and organisms, by focusing on nascent transcripts. By adapting PRO-seq and GRO-seq experimental data to analyze the mitochondrial genome we accurately identified the mtDNA TIS and transcription termination sites of both mtDNA strands in a variety of cell types and organisms, and unearthed quantitative variation in the transcription initiation of the two mtDNA strands. Additionally, we in vivo mapped, for the first time, a transcription pausing site at the light mtDNA strand of humans and other organisms. Finally, our analysis of GRO-seq and PRO-seq data that others and we generated from non-human animals enabled de-novo assembly of the entire mtDNA sequence regardless of the availability of species-specific reference genomes. Thus, our approach paves the path towards functional mtDNA genomic studies of non-model animals, far beyond RNA-seq-based studies of steady-state gene expression.

Similar to genome-wide promoters, 40-50 bp downstream to the mtDNA light strand TIS there was a read coverage peak in all human cells and most mammals. Such peaks in the nuclear genome were interpreted as RNA polymerase pausing sites. This pausing site, in human, chimpanzee, mouse and rat, exactly overlapped a known bacterial transcription pausing motif (Larson et al. 2014; Vvedenskaya et al. 2014), though in different from bacteria, this motif cannot be easily connected to translation. It is possible that the light strand pausing of the transcription machinery is required to the elongation process. Since we identified pausing only downstream to the light strand TIS, and not in the heavy strand TIS, we are inclined to interpret the pausing as functionally related to the spatially adjacent transcription-replication decision point at CSB II in human cells (Pham et al. 2006; Agaronyan et al. 2015). In consistence with this correlation, we found the mouse light strand transcription pausing site downstream to the LSP, just upstream to previously mapped RNA-DNA transition site (Chang et al. 1985). Together, these results imply that light strand transcription pausing may serve to allow sufficient time and hence enable successful transcription-to-replication transition (Fig. 5). Furthermore, the identification of TEFM, an mtDNA transcription elongation factor orthologous to the nuclear elongation factor Spt6 (Minczuk et al. 2011), suggests that mtDNA transcription pausing involves a mechanism similar to the nucleus. All this suggests that the mtDNA RNA polymerase (POLRMT) and the entire mitochondrial transcription machinery resemble the dynamics of RNA pol II.

**Figure 5.**
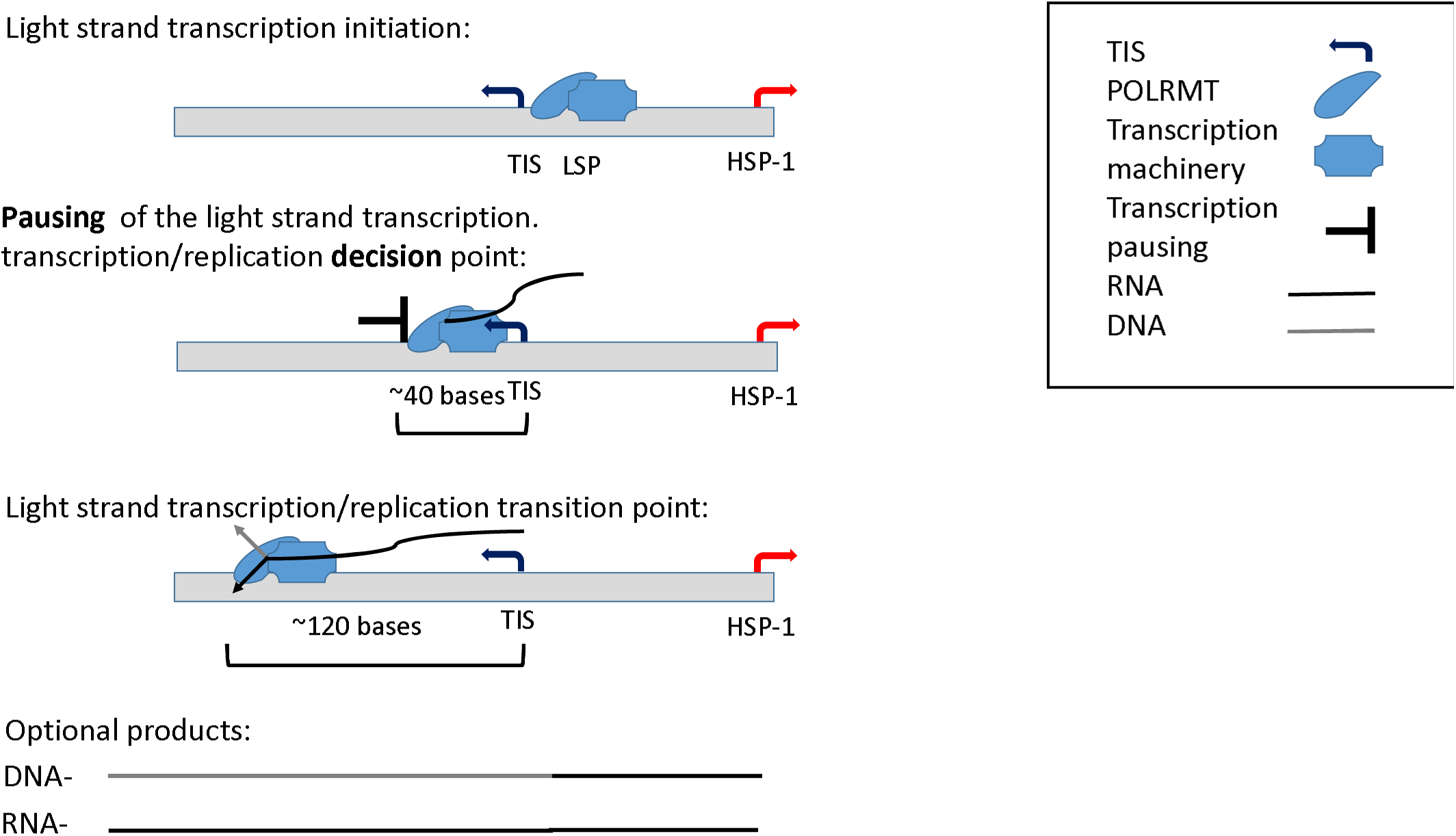
A model offering a mechanistic explanation for role of light strand transcription pausing. Grey rectangle – the mtDNA light strand.

Whereas the light strand TIS was within the same mtDNA region in all tested human cells, heavy strand TIS divided the cells into two groups: seven cell lines in which the TIS was identified within the HSP1 region (GM12004, GM12750, GM12878, HeLa, IMR90, MCF7 and U2OS) and four cell lines (K562, AC16, CD4+ and Jurkat) in which the heavy strand TIS was within the region corresponding to HSP2. Since in all 11 cell lines tested, an increase in read coverage was observed around the HSP1 region it is possible that in many cases the activity of HSP2 is masked by HSP1. Alternatively, differences in heavy strand TIS pattern among the tested cell lines may reflect the relative strength of the two heavy strand mtDNA promoters.

Analysis of a variety of human cell types revealed varying ratios between the light strand and heavy strand TIS. This may reflect differences in rates of transcription initiation, or differential proportion of pausing, in similar to findings in the nuclear genome (Kwak et al. 2013). Since these cell types also differed in their mtDNA genetic backgrounds (haplogroups), we could not determine whether the physiological differences, mtDNA sequence or even nuclear genetic variants contributed most to the observed quantitative variation in transcription initiation. As the number of analyzed samples was low, and since previous analysis of mtDNA gene expression patterns in mtDNA haplogroups revealed only subtle differences (Gomez-Duran et al. 2010; Kenney et al. 2014), association of genetic backgrounds with mtDNA TIS/pausing patterns, as well as controlled assessment of mtDNA transcription in a variety of physiological conditions, still awaits a larger sample size collected in a controlled manner.

We used our approach also to get a glimpse into the evolution of mtDNA transcription in Metazoans. We found that the pattern of mtDNA transcription (considering both initiation and termination) was very similar among the tested mammals, although quantitative differences were evident. While applying our approach to invertebrates (Drosophila and *C.elegans*), a completely different transcription pattern emerged, which correlated with the strand coding capacity. This observation raises the possibility that mtDNA gene arrangement correlated with transcription regulatory changes. This could be tested once PRO-seq/GRO-seq data becomes available from larger collection of Metazoans.

In summary, we for the first time provided accurate and quantitative analysis of mitochondrial nascent transcripts without dependence on prior sequence knowledge. We found a previously unknown evolutionarily conserved transcription pausing site downstream to the mitochondrial LSP, with likely regulatory importance for the transition between mtDNA transcription and replication. Nevertheless, we found staggering diversity in mtDNA transcription patterns among metazoans. Our approach presents previously unmatched capabilities to analyze mitochondrial transcription and assess quantitative mtDNA regulatory differences among humans, cells, physiological conditions and a variety of organisms. De novo assembly of our new data provides a means to assay the mtDNA in non-model organisms. Our findings underline the ability to measure mitochondrial transcription using the same molecular tool as is becoming more and more widely used for measuring nuclear transcription.

## Methods

### Data analyzed

CD4+ T-cell PRO-seq libraries were prepared from non-human primate and rodents species as described in established protocols [ref Danko et. al. 2015; Kwak et. al. 2013]. Blood samples (80-100mL) were obtained from three rhesus macaque and chimpanzee individuals in compliance with Cornell University IACUC guidelines. We used density gradient centrifugation to isolate peripheral blood mononuclear cells and positive selection for CD4+ cells using CD4 microbeads from Miltenyi Biotech (130-045-101 chimpanzee; 130-091-102, rhesus macaque). Mouse and rat CD4+ T-cells were isolated from splenocytes using species specific reagents from Miltenyi Biotech (130-049-201 mouse; 130-090-319 rat). In all cases enriched CD4+ T-cell nuclei were prepared by re-suspending cells in 1mL lysis buffer (10mM Tris-Cl pH 8, 300mM sucrose, 10mM NaCl, 2mM MgAc2, 3mM CaCl2, and 0.1% NP-40), washed in a wash buffer (10mM Tris-Cl pH 8, 300mM sucrose, 10mM NaCl, and 2mM MgAc2), and subsequently resuspended in 50uL of storage buffer (50mL Tris-Cl pH 8.3, 40% glycerol, 5mM MgCl2, and 0.1mM EDTA), as described (Danko et. al. 2015). For CD4+ T-cells in all species PRO-seq was performed as exactly described [kwak et. al. 2013], and sequenced using an Illumina Hi-Seq 2000 or a NextSeq 500 at the Cornell University Biotechnology Resource Center. In addition available GRO-seq and PRO-seq SRA files were downloaded from the GEO dataset. Accession numbers of each sample are listed in Supplementary Table S1. SRA files were converted into fastq format using sratoolkit (http://www.ncbi.nlm.nih.gov/Traces/sra/?view=toolkit_doc). The sequencing adaptors were trimmed by Trim-galore to reach a minimum reads length of 30 nucleotides.

### Sample-specific mtDNA sequence re-construction and mapping

As a first step to map PRO-seq and GRO-seq reads to the mtDNA, fastq files were uniquely mapped to the revised Cambridge Reference Sequence (rCRS) using BWA-aln (−q=5, −l=20, −k=2, −t=1). BWA was used to convert SAI into SAM format, which in turn was converted into a BAM file and sorted using Samtools (Li et al. 2009). Next, Samtools was used to generate VCF files of each sample (mpileup (-uf) command). Then, sample-specific mtDNA sequence was re-constructed for each of the analyzed samples using bcftools call (-c) (Samtools) in combination with vcf2fq from the vcfutils.pl package. The resulting Fastq files were uniquely re-mapped to the reconstructed sample specific mtDNA using BWA-aln (−q=5, −l=32, −k=2, −t=1), and BAM files were generated again. Removal of low MAPQ reads was performed using the Samtools ‘view’ command (−F=1804, −q=30). When analyzing non-human species we used publically available relevant mtDNA sequences (Supplementary Table S7).

### Coverage calculation

Coverage per base was calculated for a given sequence interval (separately for each strand) using Bedtolls (http://bedtools.readthedocs.org/en/latest/ version 2.25). Specifically, we employed the command ‘genomecov’ using the ‘-d’ and ‘strand’ options. For the stringent identification of pausing sites, coverage of the 3’ end of the reads was calculated using the Bedtools ‘genomecov’ command, with ‘-d’, ‘-strand’ and ‘-3’ options.

### Circular-like mapping of sample-specific mtDNA sequence

Since the mtDNA is a circular molecule and some reads may have been erroneously excluded we re-analyzed the Fastq files. To this end we remapped the reads to the sample-specific mtDNA sequence that was rearranged such, that the last 500 nucleotides of the standard mtDNA sequence were cut and pasted at the beginning of the sequence. Mapping was performed and read coverage at the former circle junction of the rearranged sequence was calculated and added to the previous mapping results.

### Pausing index

Pausing index was calculated as the ratio between the density average across 10 nucleotide sliding windows, with each position divided by the density average across the ‘gene body’. Since mtDNA genes are transcribed in polycistrones in a strand-specific manner, the ‘gene body’ was defined as the transcription unit governed by each of the strand-specific promoters. Specifically, human heavy strand ‘gene body’ was defined as the region between 100 bases downstream to the relevant identified heavy strand TIS and the end of the coding region (mtDNA position 16024). The light strand ‘gene body’ was defined as the region between mtDNA positions 3250-16024. In non-human species, ‘gene body’ was defined based on the same logic, depending on differences in the identified transcription units.

### Pausing site analysis

pausing sites were analyzed as described previously (Core et al. 2014) with some modifications. The index of each position in 200 bases downstream to the identified pausing peak was calculated. The positon having the highest pausing index score was interpreted as a candidate pausing site. Statistical significance of candidate pausing peak was calculated while assuming Poisson distribution using ‘poisson.test’ (R program), with the following parameters: ‘r’ (hypothesized rate) = the density across the ‘gene body’ and ‘alternative’ (alternative hypothesis) = ‘greater’. A peak was considered significant if p-value<0.001.

### DNAse-seq analysis

DNase-seq fastq files of the GM12878, HeLa, Jurkat, K562 and MCF7 cell lines were downloaded from the ENCODE consortium website (hgdownload-test.cse.ucsc.edu/goldenPath/hg19/encodeDCC/). DNase-seq fastq files of the CD4+ and IMR90 cell lines were downloaded from the ROADMAP Epigenomics Consortium website (http://egg2.wustl.edu/roadmap/web_portal/index.html). DNase hypersensitivity sites were identified as previously described (Blumberg et al. 2014). Briefly, for each nucleotide mtDNA position, *F*-score was calculated in sliding read windows of ~120bp using the following equation: F = (C + 1)/L + (C + 1)/R, where C represents the average number of reads in the central fragment, L represents the average reads’ count in the proximal fragment, and R represents the average reads’ count in the distal fragment. The lowest *F* scores were interpreted as a hypersensitivity site.

### NUMTs identification

NUMTs diagnosis performed in three steps: 1) BLAST screen the mtDNA (rCRS) as query in order to search NUMTs hits. 2) Collect variants which distinguish the active mtDNA from the candidate NUMTs identified in step 1. 3) Identifying and counting mtDNA mapped PRO-seq/GRO-seq reads (within BAM files) that contain NUMT variants using bam-readcount (https://github.com/genome/bam-readcount). Correlation was estimated between NUMTs variants and the nucleotide content. Since the fastq reads were trim to a length of 30 bases, BWA-aln mapability parameters were restricted to a single mismatch. The three steps of NUMTs identification was employed using two types of reference datasets: firstly, since PRO-seq and GRO-seq are based on RNA, we focused our first analysis only ‘RNA NUMTs’ and performed our initial BLAST screen against the human RefSeq RNA database. Secondly, we extended our screen to the entire human genome, utilizing the whole genome (GRCh38) and corresponding RNA as a reference.

### Assignment of sample mtDNA sequence to known genetic backgrounds – haplogroups

PRO-seq and GRO-seq sequencing reads covered a mean of 89.19% of the human heavy mtDNA strand and 85.65% of the light mtDNA strand assignment of samples to known mtDNA genetic backgrounds (haplogroups) was plausible. To this end each sample-specific mtDNA sequence was compared to the rCRS and a set of sample-specific SNPs list was generated. This data was analyzed by HaploGrep (Kloss-Brandstatter et al. 2011) and mtDNA haplogroups were assigned (Supplementary Table S8).

### De-novo mtDNA sequence assembly

We aimed at assessing whether GRO-seq and PRO-seq data was sufficient to extract the majority of the mtDNA sequence in a given species. To this end we employed CLC genomics workbench (CLC Genomics Workbench - https://www.qiagenbioinformatics.com/) to PRO-seq data from two species (Drosophila and C. elegans). Specifically, the clc_assembler command was used to de-novo assemble fastq data, employing default parameters. BWA-mem (parameters use:-B=2) was employed to map the generated contigs to the mtDNA of a phylogenetically related species that served as a scaffold (Supplementary Table S2).

### Identifying RNA-DNA differences

Previously we identified a prominent RNA-DNA difference site, common to all human samples analyzed to date (Bar-Yaacov et al. 2013). We aimed towards assessing the presence of such differences during the early stages of mtDNA tracnription. To this end BAM files generated from all tested human samples that were indexed by samtools (the index command) and analyzed by bam-readcount were used to generate metrics of nucleotide content in mtDNA nucleotide position 2617.

### Assessment of SNPs frequency

We utilized a previously published assembly of human mtDNA population variants, which stem from the analysis of nearly 10,000 individuals from diverse worldwide populations. The frequency of variants events were calculated only considering the D-Loop (mtDNA position 16024-576). The number of variants was normalized to the average of mutational events in sliding windows of 10 bases.

## Data access

We have deposited the sequencing data onto the Sequence Reads Archive (SRA), and numbers will be released upon publication.

## Acknowledgements

This study was supported by an Israeli Science foundation grant (610-12) awarded to D.M., Binational-Science Foundation grant (2013060) awarded to D.M. and A.K, and a National Heart, Lung, and Blood Institute grant (UHL129958A) to CGD. The authors also would like to thank the Harbor foundation for a scholarship for excellent PhD students awarded to A.B.

## Supplementary figures

**Supplementary Figure S1. NUMTs from *Homo sapiens* Ref Seq RNA database**.

**Supplementary Figure S2. De-novo assembly of invertebrates species**. PRO-seq and GRO-seq data enabled de-novo assembly of the entire invertebrate mtDNA sequence without a reference sequence. Left panel – C. elegans. Right panel – Drosophila. Scaffold used for de novo assembly: for Drosophila – *Bactrocera arecae* mtDNA and for C. elegans - mtDNA from *Litoditis aff. Marina Pml*. Blue cycle – mapped de-novo assembly contigs. Black cycle – the scaffold mtDNA sequence used to map the de novo assembly contigs. For C. elegans, the mtDNA reads covered 97% of the scaffold genome, and for Drosophila they covered 80% of the scaffold genome. Notably, the gaps mostly correspond to the non-coding control region.

**Supplementary Table S1**. SRA accession numbers of samples used in this study.

**Supplementary Table S2**. Nucleotide positions with mutations that correspond to NUMTs.

**Supplementary Table S3**. Summary statistics of the sequence mapping process.

**Supplementary Table S4**. Calculated values of Human mtDNA pausing site density.

**Supplementary Table S5**. RNA-DNA Difference (RDD) in mtDNA position 2617.

**Supplementary Table S6**. Ratio of lights strand coverage versus heavy strand coverage across mammals.

**Supplementary Table S7**. List of mitochondrial reference genomes used in this study.

**Supplementary Table S8**. Samples haplogroups.

